# Inhibition of mitochondrial complex III causes selective dopaminergic neurotoxicity by redox stress in *Caenorhabditis elegans*

**DOI:** 10.1101/2025.10.21.683798

**Authors:** Javier Huayta, Aliyah S. Webster, Jay Jhaver, Joel N. Meyer

## Abstract

Environmental factors including chemical exposures are important contributors to Parkinson’s disease (PD). Nearly all well-validated chemicals involved in PD affect mitochondria, and the great majority of those identified inhibit mitochondrial complex I, causing ATP depletion and oxidative stress. We hypothesized that inhibition of mitochondrial complex III would also cause dopaminergic neurotoxicity. Using *Caenorhabditis elegans* to evaluate the *in vivo* effects of complex III-inhibiting pesticides antimycin A and pyraclostrobin, we found that both caused selective dopaminergic neurotoxicity. We evaluated exacerbation of dopaminergic neurotoxicity by the presence of α-synuclein, and *pdr-1*/PRKN and *pink-1*/PINK1 mutant backgrounds and found increased neurotoxicity for *pdr-1*. Complex III inhibition caused a more-oxidized cellular environment in those neurons and pharmacological and genetic antioxidant interventions rescued neurotoxicity, but energetic rescue attempts did not. Finally, optogenetic production of superoxide anion specifically at complex III caused dopaminergic neuronal damage. Thus, redox stress at complex III following chemical exposure causes dopaminergic neurotoxicity *in vivo* in *C. elegans*.

## Introduction

Parkinson’s disease (PD), characterized by loss of dopaminergic neurons in the *substantia nigra*, affected 10 million people globally in 2021 [1] and is projected to increase to 25.2 million by 2050 [2]. This increase is the outcome of an increasingly older population worldwide, particularly in the most polluted parts of the world [3]. Environmental factors are important contributors to PD, and many studies have demonstrated the role of chemical exposures [4,5]. Most if not all the chemicals linked to onset of PD affect mitochondria [6,7]. However, there is strong evidence for association with PD for only a few chemicals, and because relatively few people are exposed to significant amounts of those chemicals, they collectively likely explain only a small fraction of idiopathic PD. It is not feasible to comprehensively test all the chemicals that induce mitochondrial dysfunction for involvement in PD or parkinsonism, as these number in the hundreds to thousands [8]. However, mitochondrial dysfunction is a broad term that includes many specific mechanisms of toxicity [9–11]. These include inhibition of all four electron transport chain complexes, ATP synthase, and Krebs cycle enzymes; redox cycling; mitochondrial DNA damage; and uncoupling of ATP production from oxygen consumption. Therefore, to prioritize chemicals for testing, it would be valuable to identify which specific types of mitochondrial toxicity drive dopaminergic neurotoxicity. To date, there is limited evidence for the role of specific types of mitochondrial dysfunction in dopaminergic neurotoxicity. There is strong evidence for complex I inhibition [12,13], and one report that mitochondrial uncoupling may be protective against dopaminergic neurodegeneration [14]. A study using cell culture models examined 21 different pesticides that inhibited complex I, II, or III, finding evidence that some inhibitors of complex I and complex III, but no complex II inhibitors, were selectively toxic to dopaminergic cells in culture [15].

One reason to hypothesize that not all forms of mitochondrial dysfunction cause dopaminergic neurotoxicity is that these forms of mitochondrial dysfunction do not all cause the same downstream molecular events. Two common downstream outcomes of mitochondrial toxicity are redox imbalance and ATP depletion [16,17], but these key events will not universally result from all forms of mitochondrial toxicity. For example, complex I inhibitors will typically both decrease ATP availability and increase reactive oxygen species (ROS) by diverting electron flow from ubiquinone to molecular oxygen [12], while a mitochondrial uncoupler may also decrease ATP levels but decrease ROS by decreasing mitochondrial membrane potential (MMP) [14]. There is some literature addressing the importance of redox imbalance in comparison to ATP depletion in mitochondrial toxicant-induced dopaminergic neurotoxicity. For example, two studies indicate that although the complex I inhibitor rotenone both decreased ATP levels and increased ROS, it was ROS that drove neurodegeneration [12,18]. However, careful mechanistic analysis of this sort after chemical exposure has been rare. It is important to determine which forms of mitochondrial dysfunction are important for both regulatory and mechanistic reasons.

In this work, we use the model organism *C. elegans*, a well-characterized model for neurotoxicity [19], to evaluate the *in vivo* effects of complex III inhibition on dopaminergic neurotoxicity, ATP levels, and redox state. To ensure that results are physiologically relevant, informative of potential selectivity for dopaminergic neurotoxicity, and not simply part of a generalized toxic response, we use exposure concentrations that cause mild and no organismal-level toxicity, and compare outcomes in dopaminergic neurons to several other neuron types present in *C. elegans*. We also evaluate potential exacerbation of dopaminergic neurotoxicity by presence of α-synuclein and genetic mitophagy disruption. We demonstrate that developmental exposure *in vivo* to pesticides antimycin A and pyraclostrobin leads to dopaminergic neurotoxicity. Finally, using a variety of pharmacological, genetic, and optogenetic tools, we show that this neurotoxicity is caused by increased mitochondrial ROS at complex III.

## Results

### Pesticides that inhibit mitochondrial complex III cause selective dopaminergic neurotoxicity *in vivo*, in the absence of obvious generalized organismal toxicity

We first asked whether exposure to complex III inhibitors would cause selective dopaminergic neurotoxicity. We took two steps to ensure that we were measuring a selective outcome: 1) we identified concentrations that caused relatively mild organismal-level toxicity (decreased larval growth), and 2) we compared effects in dopaminergic neurons to all other populations of neurons.

Exposure to mitochondrial toxicants causes growth inhibition in *C. elegans* [20,21]. We quantified the effects on *C. elegans* growth of developmental exposure to two inhibitors of coenzyme Q – cytochrome *c* oxidoreductase (complex III) of the ETC: antimycin A, a piscicide that binds to the ubiquinone Qi site [22]; and pyraclostrobin, a fungicide that binds to the ubiquinone Q0 site [23] (Fig. 1A). We observed a clear decrease in average worm length with increasing antimycin A concentration (Fig. 1B) but could not identify a significant decrease in average worm length after exposure to concentrations of pyraclostrobin up to 50 µM, the limit of solubility (Fig. 1C). We used this data to construct dose response curves [24] and calculated the concentrations causing a reduction of 10% and 25% in average worm length for antimycin A to be 100 and 500 nM respectively (Fig. S1A). We used these EC_10_ and EC_25_ values for subsequent experiments to ensure that our neuronal endpoint measurements were not secondary to generalized whole organism toxicity. Because we could not test pyraclostrobin over 50 µM, we assigned this maximum concentration as our EC_10_ value (Fig. S1B). Therefore, for further experiments, we used these concentrations in addition to low concentrations of 10 nM for antimycin A and 10 µM for pyraclostrobin.

**Figure 1.**
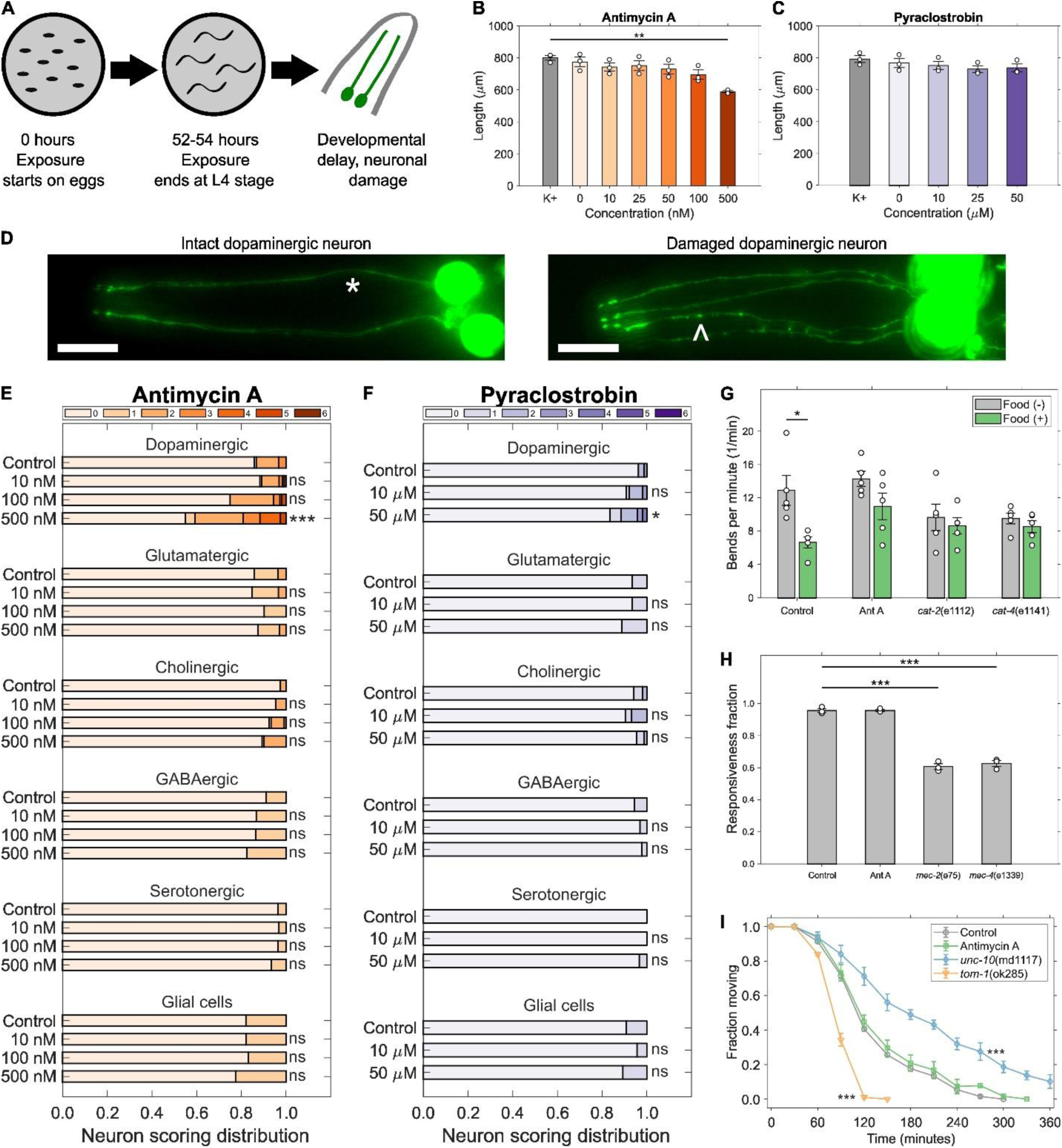
Mitochondrial complex III inhibition causes selective dopaminergic neuronal damage. (**A**) Chemical developmental exposure occurs from the embryo to the L4 stage; after 52-54 hours endpoint measurement was performed. Length of worms exposed to increasing concentrations of (**B**) antimycin A and (**C**) pyraclostrobin during development. *N* = 3 biological replicates, *n* = 50 – 100 worms per treatment per replicate, error bars SEM, one-way ANOVA with Dunnet’s post-hoc test compared to 0 nM or 0 µM controls (K+ is a control without DMSO vehicle), \*\**P* < 0.01. (**D**) Representative images of intact (*) and damaged neurites (^) of dopamine neurons. Neuronal damage scoring distribution for dopaminergic, glutamatergic, cholinergic, GABAergic, and serotonergic neurons, and glial cells located in the head of worms developmentally exposed to (**E**) antimycin A and (**F**) pyraclostrobin. *N* = 3 biological replicates, *n* = 40 dendrites for dopaminergic, glutamatergic, and cholinergic neurons, and glial cells, 30 dendrites for GABAergic neurons, 20 dendrites for serotonergic neurons, per treatment per replicate, chi-square test with Bonferroni post-hoc test, \**P* < 0.05, \*\*\**P* < 0.001. (**G**) Basal slowing response, (**H**) sensitivity to soft touch, and (**I**) aldicarb-induced paralysis after developmental exposure to antimycin A. *N* = 5 biological replicates, *n* = 10 worms per treatment per replicate for (G), *N* = 3 biological replicates, *n* = 20 - 30 worms per treatment per replicate for (H) and (I), error bars SEM, two-way ANOVA with Tukey’s HSD post-hoc test for (G), one-way ANOVA with Dunnett’s post-hoc test for (H), log-rank test for (I), \**P* < 0.05, \*\*\**P* < 0.001.

Next, we tested whether these exposures, causing mild or no growth impairment, would cause neurotoxicity. Diminished mitochondrial function is linked to increased neurodegeneration [9], and neurons show strong sensitivity to mitochondrial impairment [10,11]. However, different toxicants generally affect different types of neurons [25]. Therefore, we quantified neuron morphological damage in multiple neuronal types, quantifying morphological irregularities including kinks, blebs, and loss of continuity in dendrites located in the head region (Fig. 1D, S2) using previously described scoring systems [21,26]. Developmental exposure to antimycin A led to a statistically significant increase in dopaminergic neuronal damage (Fig. 1E). However, we did not find significant degeneration after exposure to antimycin A in glutamatergic, cholinergic, GABAergic, or serotonergic neurons, or glial cells (Fig. 1E). Similarly, developmental exposure to pyraclostrobin caused a statistically significant increase in dopaminergic neuron damage (Fig. 1F), albeit not as substantial as that caused by 500 nM antimycin A. Comparable to the results with antimycin A, no concentration of pyraclostrobin induced neurotoxicity in other neuron types or glial cells (Fig. 1F).

To further test the selectivity of these effects for dopamine neurons, we tested behaviors and phenotypes linked to neuronal function. First, we evaluated the basal slowing response (BSR), a food-scavenging behavior regulated by dopaminergic sensory neurons that causes worms to slow down in the presence of bacterial food [27], with loss of BSR resulting from impaired dopaminergic function [28]. We found that developmental exposure to 500 nM antimycin A abrogated the BSR of worms on plates with food (Fig. 1G). We used *cat-2* and *cat-4* mutant strains that are defective for BSR as positive controls. We also tested worms exposed during development to 500 nM antimycin A for their sensitivity to soft touch and for their rate of paralysis in the presence of the nematicide aldicarb. Exposed worms did not lose their recoiling behavior as response to touch in their heads and tails (Fig. 1H), a behavior controlled by mechanosensory glutamatergic neurons [29]. Similarly, exposed worms did not diverge from controls in their rate of paralysis in the presence of aldicarb (Fig. 1I), a phenotype regulated by acetylcholine synthesis and release [30]. As positive controls for these assays, we used *mec-2* and *mec-4* mutant strains defective for soft touch and *tom-1* and *unc-10* mutant strains sensitive and resistant to aldicarb, respectively.

### Deletion of mitophagy gene *pdr-1*, but not *pink-1* or the presence of α-synuclein, increases developmental neurotoxicity driven by complex III inhibition

We hypothesized that dopaminergic neurotoxicity induced by inhibition of complex III would be further increased by the presence of other stressors, considering evidence that PD is often the result of a combination of environmental and genetic factors [31]. The protein α-synuclein, encoded by the SNCA gene, is implicated in the onset of PD and other human neurodegenerative diseases [31,32]. In these conditions, α-synuclein misfolds and aggregates in Lewy bodies. This α-synuclein buildup can damage neurons and cause symptoms of these neurodegenerative diseases [32]. We tested if the presence of α-synuclein would alter the effects of complex III inhibitors. Because *C. elegans* does not produce endogenous α-synuclein, we used a transgenic *C. elegans* strain containing the human SNCA gene expressed specifically in dopaminergic neurons [33]. We also tested the effect of mutations in *pdr-1* and *pink-1*, the *C. elegans* homologs of the human PRKN and PINK1 Parkinson’s disease genes, respectively. They are involved in mitophagy, and their loss leads to impaired mitochondrial function and PD [34].

The expression of α-synuclein alone had a large effect on dopaminergic neurons at all concentrations of antimycin A and pyraclostrobin tested, including controls (Fig. 2A-B). The high concentration of 500 nM antimycin A still had a significant effect on dopaminergic neurotoxicity in the presence of α-synuclein, and the *pdr-1* and *pink-1* mutations (Fig. 2A). No significant differences in neurotoxicity were found when exposing worms with the presence of both α-synuclein and either *pdr-1* or *pink-1* backgrounds to antimycin A (Fig. 2A). The neurotoxic effect on dopamine neurons observed with 50 µM pyraclostrobin was recapitulated only for worms with the *pdr-1* background, but not for those with the α-synuclein or *pink-1* backgrounds (Fig. 2B). Exposure to pyraclostrobin in combination with the presence of α-synuclein and either *pdr-1* or *pink-1* mutations did not cause significant differences in neuronal damage.

**Figure 2.**
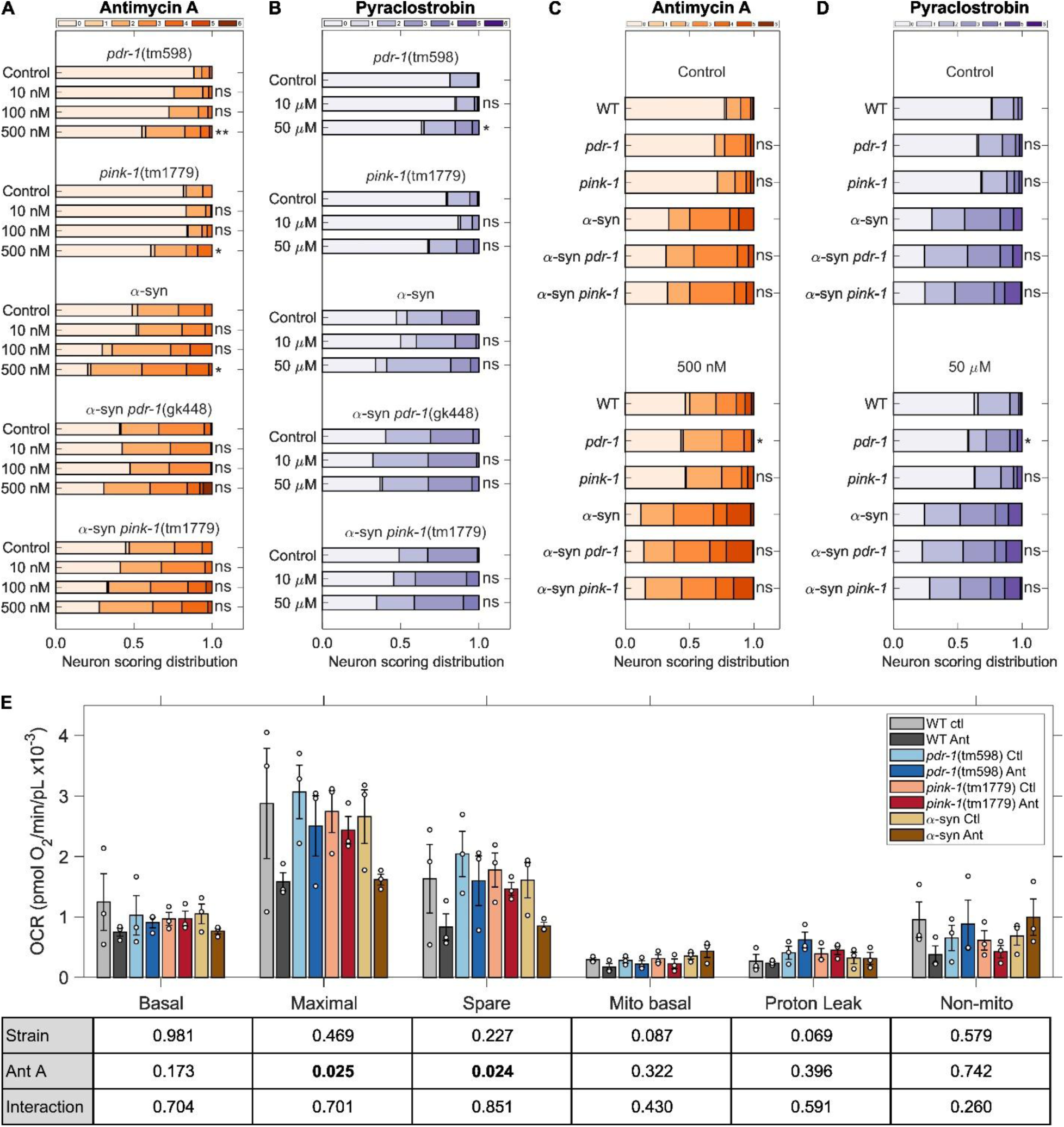
Loss of mitophagy gene *pdr-1* independently regulates neurotoxicity after developmental inhibition of complex III. Neuronal damage scoring distribution for cephalic dopaminergic neurons in combination with dopaminergic α-synuclein, a *pdr-1* mutant background, a *pink-1* mutant background, dopaminergic α-synuclein and *pdr-1* mutant background, and dopaminergic α-synuclein and *pink-1* mutant background in worms developmentally exposed to increasing concentrations of (**A**) antimycin A and (**B**) pyraclostrobin. *N* = 3 biological replicates, *n* = 40 dendrites per treatment per replicate, chi-square test with Bonferroni post-hoc test, ns *P* > 0.05, \**P* < 0.05, \*\**P* < 0.01. Neuronal damage scoring distribution for cephalic dopaminergic neurons to compare the effects of genetic background after developmental exposure to (**C**) 500 nM antimycin A and (**D**) 50 µM pyraclostrobin. *N* = 3 biological replicates, *n* = 60 - 80 dendrites per treatment per replicate, chi-square test with Bonferroni post-hoc test, ns *P* > 0.05, \**P* < 0.05. (**E**) Oxygen consumption rate of wild type, *pdr-1* mutant, *pink-1* mutant, and α-synuclein carrying worms after developmental exposure to antimycin A. *N* = 3 biological replicates, *n* = 20 – 30 worms in each of 5 technical replicates, error bars SEM, two-way ANOVA with Tukey’s HSD post-hoc test for each OCR category, bold *P* values in table are < 0.05.

To directly compare effects of antimycin A and pyraclostrobin in different strains including wildtype, we carried out a second set of exposures (Fig. 2C-D). Because concentrations lower than 500nM antimycin A or 50 µM pyraclostrobin did not cause increased dopaminergic neurotoxicity in the presence of α-synuclein and/or *pdr-1* and *pink-1* mutant backgrounds, we evaluated the neurotoxic effects driven by differences between strains only at these highest concentrations. We observed no significant difference between the *pdr-1* and *pink-1* strains compared to the wild type for non-exposed worms (Fig. 2C-D). The same result was observed for the *pdr-1* and *pink-1* strains with α-synuclein presence compared to the α-synuclein wild type: the *pdr-1* and *pink-1* deletions did not further exacerbate neurotoxicity caused by α-synuclein expression (Fig. 2C-D). This last result was repeated for worms exposed to antimycin A and pyraclostrobin in the presence of α-synuclein (Fig. 2C-D). Worms developmentally exposed to antimycin A or pyraclostrobin with a *pdr-1* mutant background showed a significant increase in dopaminergic neuronal damage with respect to the exposed wild type worms (Fig. 2C-D). This result was not observed for the *pink-1* mutant background.

Next, we explored if these results could be driven by increased mitochondrial dysfunction driven by these interactions. We measured the oxygen consumption rate (OCR) of wild type, *pdr-1* mutant, *pink-1* mutant, and α-synuclein worms after developmental exposure to 500 nM antimycin A (Fig. 2E). Oxygen consumption is an integral part of oxidative phosphorylation (OXPHOS) in mitochondria, with changes in mitochondrial respiration indicating disruption of mitochondrial function [35]. Under these conditions, we found no effects of antimycin A exposure or strain differences on basal OCR, mitochondrial basal OCR, proton leak or non-mitochondrial OCR (Fig. 2E). Mitochondrial basal OCR is the amount of oxygen consumed to convert ADP to ATP; proton leak indicates the oxygen consumption in mitochondrial after ATP synthase is inhibited (i.e., OCR consumed by mitochondria, but for which the resulting proton gradient is used for purposes other than ATP synthesis); and non-mitochondrial OCR is a measure of the cellular processes that consume oxygen outside of the electron transport chain. In contrast, antimycin A exposure but not strain differences significantly influenced maximal OCR and spare capacity (Fig. 2E). Maximal OCR reflects the amount of oxygen consumed after chemically uncoupling mitochondria, causing a maximal rate of consumption; the spare capacity is the difference between maximal and basal OCR. However, post-hoc analysis for comparisons within maximal OCR and spare capacity did not reveal significant differences between treated and untreated worms of the same strain.

Finally, we considered that developmental inhibition of complex III could sensitize dopaminergic neurons to late life neurodegeneration, with or without further chemical insults. To test this, we allowed developmentally exposed worms to reach day 6 of adulthood, and we quantified dopaminergic neurodegeneration at this point. We found no significant differences between worms exposed to antimycin A during development and controls (Fig. S3). In addition, we exposed worms developmentally exposed to antimycin A to secondary challenge with the well-characterized dopaminergic neurotoxin 6-hydroxydopamine (6-OHDA) [36]. Worms exposed to antimycin A followed by 6-OHDA challenge show a significant increase in dopaminergic neurotoxicity compared to control worms exposed only to 6-OHDA (Fig. S4).

### Complex III inhibition generates an oxidized redox state in mitochondria of dopaminergic neurons

Inhibition of complex III may decrease the ATP:ADP ratio as electron transfer and proton pumping are inhibited, at least if the exposure is high enough. It may also lead to generation of ROS as electrons leak to oxygen [37,38]. However, there is limited evidence for *in vivo* impacts of complex III inhibition, in particular in specific cells. Furthermore, these effects are dose - dependent, raising the question of whether they occur in our experimental conditions. Therefore, we characterized how our chemical exposures altered bioenergetic and redox balance. We used 500 nM antimycin A and 50 µM pyraclostrobin because these were the concentrations that induced significant dopaminergic neurotoxicity after developmental exposure. We evaluated the redox state in dopaminergic neurons by using a roGFP reporter under the *dat-1* promoter and localized to mitochondria [39,40]. Similarly, we quantified the ATP:ADP ratios of mitochondria in dopaminergic neurons by using the PercevalHR reporter under the *dat-1* promoter [40,41].

We found that after developmental inhibition of complex III (Fig. 3A), neither antimycin A nor pyraclostrobin caused an increase in oxidized:reduced roGFP (Fig. 3B, S5A). We used a short-term exposure to the dopaminergic neurotoxin 6-OHDA, previously reported to induce an increase in oxidized:reduced roGFP [12,40], as a positive control. Similarly, neither developmental exposure to antimycin A nor pyraclostrobin led to significant changes in the ATP:ADP ratio in dopaminergic neurons (Fig. 3C, S5B). We also used 6-OHDA as a positive control for these tests. We considered that the 52 to 54 hours duration of developmental exposure may have enabled the worms to produce compensatory mechanisms to ameliorate mitochondrial ROS generation [42,43] or energetic challenge [44,45]. To test this, we repeated the developmental exposures to 500 nM antimycin A but measured redox and bioenergetic changes in dopaminergic neurons after 18 and 24 hours (Fig. 3D). We found that 18 hours after starting exposure, antimycin A caused a significant increase in oxidized:reduced roGFP (Fig. 3E, S5C). However, this effect was no longer significant at the 24-hour timepoint. Notably, no significant changes in ATP:ADP ratios were measured for either the 18- or 24-hour timepoints (Fig. 3F, S5D). In addition, we tested an acute exposure to a high concentration of 10 µM antimycin A (20 times the high concentration for developmental exposure of 500 nM) for 2.5 hours on L4 stage worms (Fig. 3G). This short exposure led to a significant increase in oxidized:reduced roGFP (Fig. 3H, S5E), but no significant change in ATP:ADP ratio was measured (Fig. 3I, S5F). Worms affected by this acute exposure to antimycin A did not show significant neuronal damage immediately after exposure (Fig. S6), suggesting that this exposure was too short to enact the neurotoxic effects of antimycin A.

**Figure 3.**
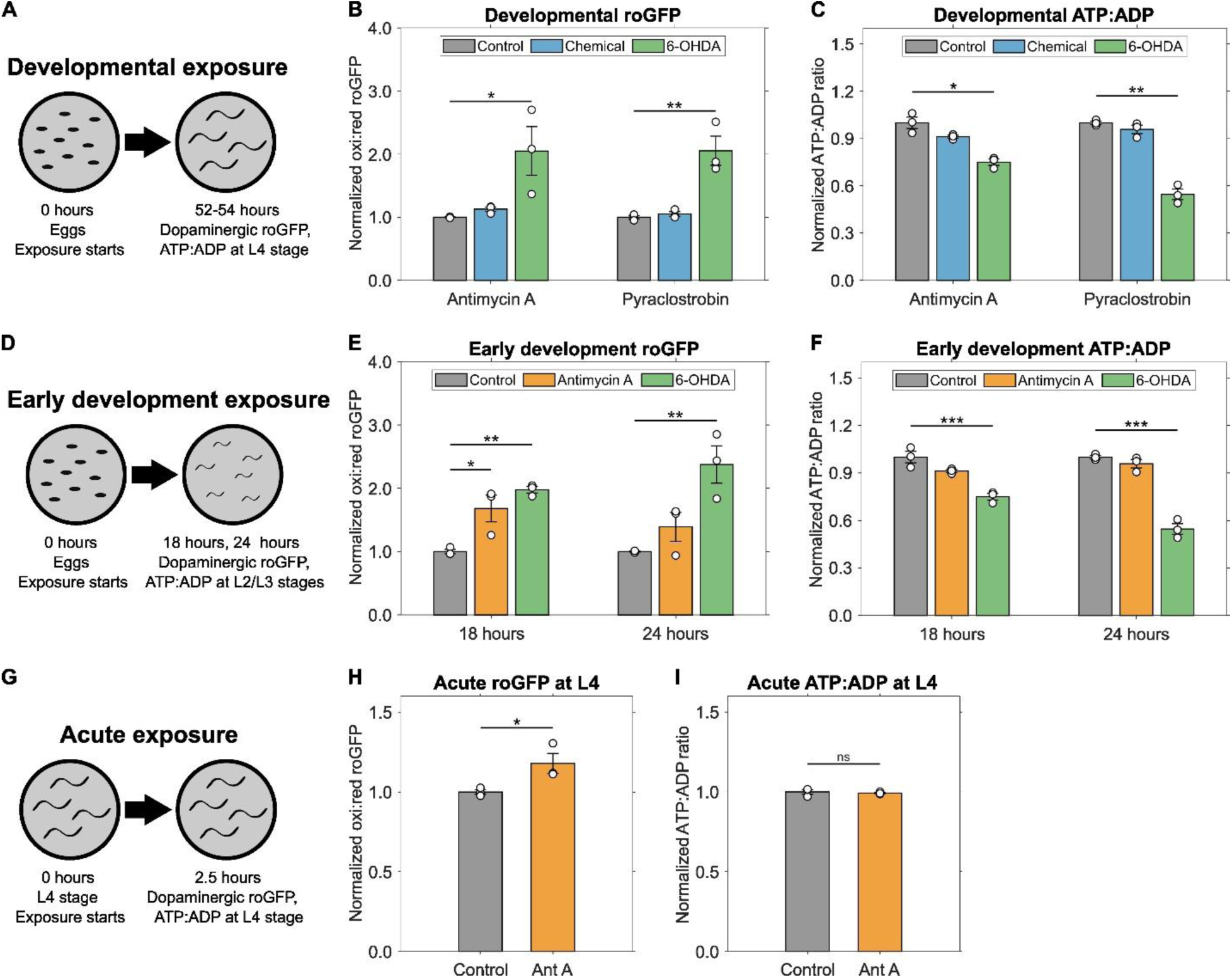
Complex III inhibition alters dopaminergic mitochondrial redox but not bioenergetic state. (**A**) Schematic of developmental exposure. (**B**) Oxidized to reduced roGFP ratio in cephalic dopaminergic neurons of *C. elegans* developmentally exposed to 500 nM antimycin A and 50 µM pyraclostrobin. (**C**) ATP to ADP ratio as measured using the PercevalHR reporter in cephalic dopaminergic neurons of *C. elegans* developmentally exposed to 500 nM antimycin A and 50 µM pyraclostrobin. 6-OHDA included as positive control. *N* = 3 biological replicates, *n* = 20 neurons per treatment per replicate, error bars SEM, one-way ANOVA test with Dunnett’s post-hoc test for each chemical, \**P* < 0.05, \*\**P* < 0.01. (**D**) Schematic of early development exposure. (**E**) Oxidized to reduced roGFP ratio in cephalic dopaminergic neurons of *C. elegans* exposed to 500 nM antimycin for 18 and 24 hours. (**F**) ATP to ADP ratio as measured using the PercevalHR reporter in cephalic dopaminergic neurons of *C. elegans* exposed to 500 nM antimycin for 18 and 24 hours. 6-OHDA included as positive control. *N* = 3 biological replicates, *n* = 40 neurons per treatment per replicate, error bars SEM, one-way ANOVA test with Dunnett’s post-hoc test for each time point, \**P* < 0.05, \*\**P* < 0.01, \*\*\**P* < 0.001. (**G**) Schematic of acute exposure. (**H**) Oxidized to reduced roGFP ratio and (**I**) ATP to ADP ratio as measured using the PercevalHR reporter in cephalic dopaminergic neurons of *C. elegans* exposed to 10 µM antimycin A for 2.5 hours at the L4 larval stage. *N* = 3 biological replicates, *n* = 20 neurons per treatment per replicate, one-way ANOVA test with Dunnett’s post-hoc test, ns *P* > 0.05, \**P* < 0.05. All roGFP and ATP:ADP values normalized to their respective controls.

### S3QEL-2 rescues neurotoxicity caused by complex III inhibition

We next directly tested whether neurotoxicity induced by inhibition of complex III is caused by redox or bioenergetic stress by performing pharmacological co-exposures. N-acetylcysteine (NAC) is a well-characterized antioxidant that works by several mechanisms: it is a glutathione precursor; it directly reacts with free radicals through its thiol group; it acts as reducing agent that can break disulfide bonds in inappropriately oxidized proteins; and it promotes the production of intracellular hydrogen sulfide that also serves as an antioxidant [46–48]. We developmentally exposed *C. elegans* to 500 nM antimycin A in conjunction with 2.5 mM NAC (Fig. 4A). NAC by itself did not have a significant effect on neurotoxicity, and an apparent decrease in antimycin A-mediated neurotoxicity with NAC co-exposure did not reach statistical significance. We next tested S3QEL-2, a selective inhibitor of superoxide anion production from the outer Q-binding site of complex III that acts without affecting OXPHOS [49,50]. Exposure to 100 µM S3QEL-2 alone did not cause detectable neurotoxicity, but rescued antimycin A-mediated neurotoxicity (Fig. 4A). Thus, blocking ROS production at complex III with S3QEL-2 was sufficient to offset the neurotoxic effects of antimycin A-mediated complex III inhibition.

**Figure 4.**
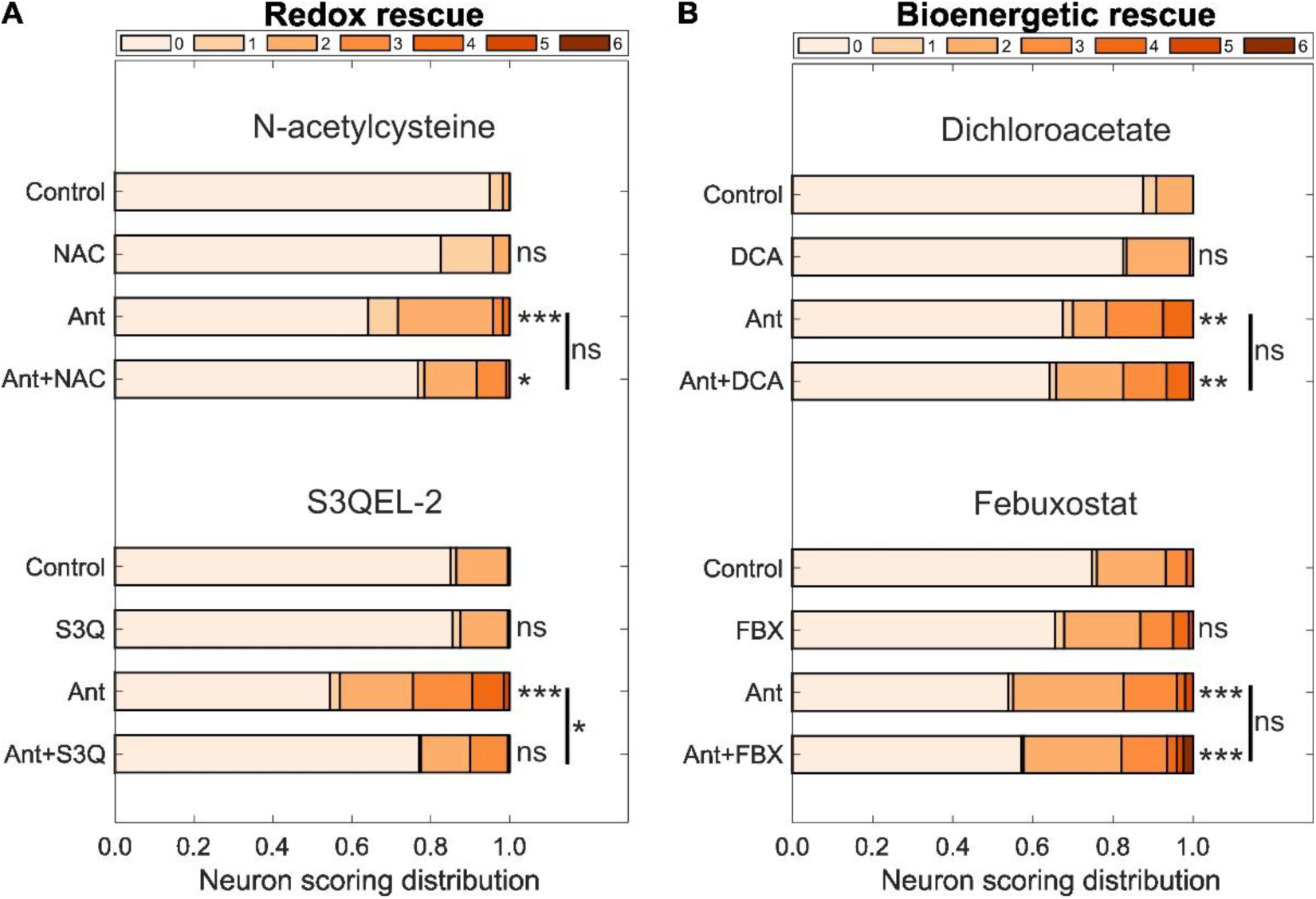
Antioxidant rescue of neurotoxicity after developmental inhibition of complex III. (**A**) Neuronal damage scoring distribution for cephalic dopaminergic neurons after developmental exposure to 500 nM antimycin A in combination with 2.5 mM N-acetylcysteine and 100 µM S3QEL-2. (**B**) Neuronal damage scoring distribution for cephalic dopaminergic neurons after developmental exposure to 500 nM antimycin A in combination with 25 mM dichloroacetate and 10 µg/mL febuxostat. *N* = 3 biological replicates (4 biological replicates for S3QEL-2), *n* = 40 - 60 dendrites per treatment per replicate, chi-square test with Bonferroni post-hoc test, ns *P* > 0.05, \**P* < 0.05, \*\**P* < 0.01, \*\*\**P* < 0.001.

To test bioenergetic rescue, we first tested dichloroacetate (DCA), a synthetic organic acid that inhibits pyruvate dehydrogenase kinase, which inhibits pyruvate dehydrogenase [46,51]. DCA thus enhances the activity of pyruvate dehydrogenase, which in turn converts pyruvate into acetyl-CoA that is used for ATP production in mitochondria [52,53]. By diverting pyruvate away from less-energetically favorable glycolysis, DCA treatment may provide a bioenergetic rescue. Developmental exposure to 25 mM dichloroacetate did not reduce the neurotoxic effect of antimycin A on dopaminergic neurons (Fig. 4B). We next tested febuxostat (FBX), which blocks the activity of xanthine oxidase, an enzyme responsible for converting purines to uric acid, which can support the purine salvage pathway and ATP regeneration. FBX has been shown to increase ATP levels and protect mitochondria in a *C. elegans* model of muscle deterioration [54].

Developmental exposure to 10µg/mL FBX did not alter neurotoxicity in dopaminergic neurons and did not affect the neurotoxic effects of antimycin A (Fig. 4B). Overall, our pharmacological efforts to increase ATP availability to offset any potential bioenergetic effects caused by antimycin A did not alter its dopaminergic neurotoxicity.

We also evaluated the effects of all four chemicals on dopaminergic mitochondrial redox and bioenergetics states. Developmental exposure to 2.5 mM NAC, 100 µM S3QEL-2, and 25 mM DCA did not cause significant changes in oxidized:reduced roGFP, either by themselves or in combination with antimycin A (Fig. S7A). For S3QEL-2, developmental exposure to 500 nM antimycin A caused a significant increase in oxidized:reduced roGFP, but 100µM S3QEL-2 rescued 500 nM antimycin A-induced oxidation (Fig. S7A). Similarly, neither developmental exposure to 2.5 mM NAC, 100 µM S3QEL-2, 25 mM DCA, 500 nM antimycin A, or their combinations produced significant changes in ATP:ADP ratios in dopaminergic neurons (Fig. S7B). We also tested whether DCA or FBX affect ATP levels when measured during early development but did not observe a significant effect (Fig. S7C; the trends toward decreased ATP after antimycin A, apparently rescued by DCA and FBX, were not significant).

Finally, we evaluated rescue with myxothiazol, because this chemical has been reported to inhibit the formation and release of semiquinone radicals in complex III [55,56], indicating a potential opposite effect to antimycin A. We tested the effects of 10 µM myxothiazol along with 500 nM antimycin A. Myxothiazol did not reduce the neurotoxic effects of antimycin A (Fig. S7D). It is possible that this is the result of substrate-dependent enhancement of the release of hydrogen superoxide in complex III or a concentration-dependent effect [57]. In a similar result to the other chemicals tested, we did not find an effect of myxothiazol on dopaminergic oxidized:reduced roGFP or ATP:ADP ratios (Fig. S7E).

### Mitochondrial superoxide anion regulates neurotoxicity caused by complex III inhibition

S3QEL-2 is described as a specific inhibitor of superoxide anion production in complex III [49], but most characterization of S3QEL-2 effects has been *in vitro*. To validate our S3QEL-2-based mechanistic evidence for a causative role for superoxide anion production, we manipulated mitochondrial manganese superoxide dismutase, which has a primary role in scavenging superoxide anion radicals in mitochondria [58]. *C. elegans* carries two mitochondrial superoxide dismutase genes, *sod-2* and *sod-3* [59]. Loss-of-function mutations in *sod-2* or *sod-3* did not dramatically increase dopaminergic neurotoxicity on their own or after antimycin A exposure (Fig. 5A). However, it is possible that complementary mechanisms that target mitochondrial ROS are sufficient to ameliorate its effects [42]. With this consideration, we evaluated if increasing superoxide dismutase availability would influence neurotoxicity. Using a strain containing both the dopaminergic reporter and multiple copies of *sod-2* causing overexpression [60], we tested the effects of complex III inhibition. Increasing expression of *sod-2* abrogated the neurotoxic effects of exposure to 500 nM antimycin A (Fig. 5B). As a second test of whether supplementation of superoxide dismutase activity reduces dopaminergic neurotoxicity, we performed chemical rescue with the superoxide dismutase and catalase mimetic EUK-134 [61]. This synthetic antioxidant converts superoxide anion into hydrogen peroxide and then converts hydrogen peroxide into water and oxygen [62]. We found that the addition of 0.5 mM EUK-134 during development to worms exposed to 500 nM antimycin A significantly reduced the dopaminergic neurotoxicity caused by antimycin A itself (Fig. 5B).With three lines of evidence that complex III inhibitors cause dopaminergic neurotoxicity via production of superoxide anion, we finally asked whether artificial production of superoxide anion specifically at complex III would cause dopaminergic neurotoxicity. We tested *C. elegans* containing the dopaminergic reporter and SuperNova appended to *ucr-2.3* [63], a gene encoding a complex III core protein located in the mitochondrial matrix [64]. SuperNova is an optogenetically activated protein that upon excitation produces ROS, particularly superoxide anion and singlet oxygen [65]. This chromophore enables spatially targeted generation of these ROS at complex III, localized to the mitochondrial matrix [63]. We found that worms with light-activated SuperNova during development show significant damage in dopaminergic neurons when compared to controls (Fig. 5C). This confirms, in a manner independent from chemical exposure, that ROS generation at the complex III site causes dopaminergic neuronal damage in our model.

**Figure 5.**
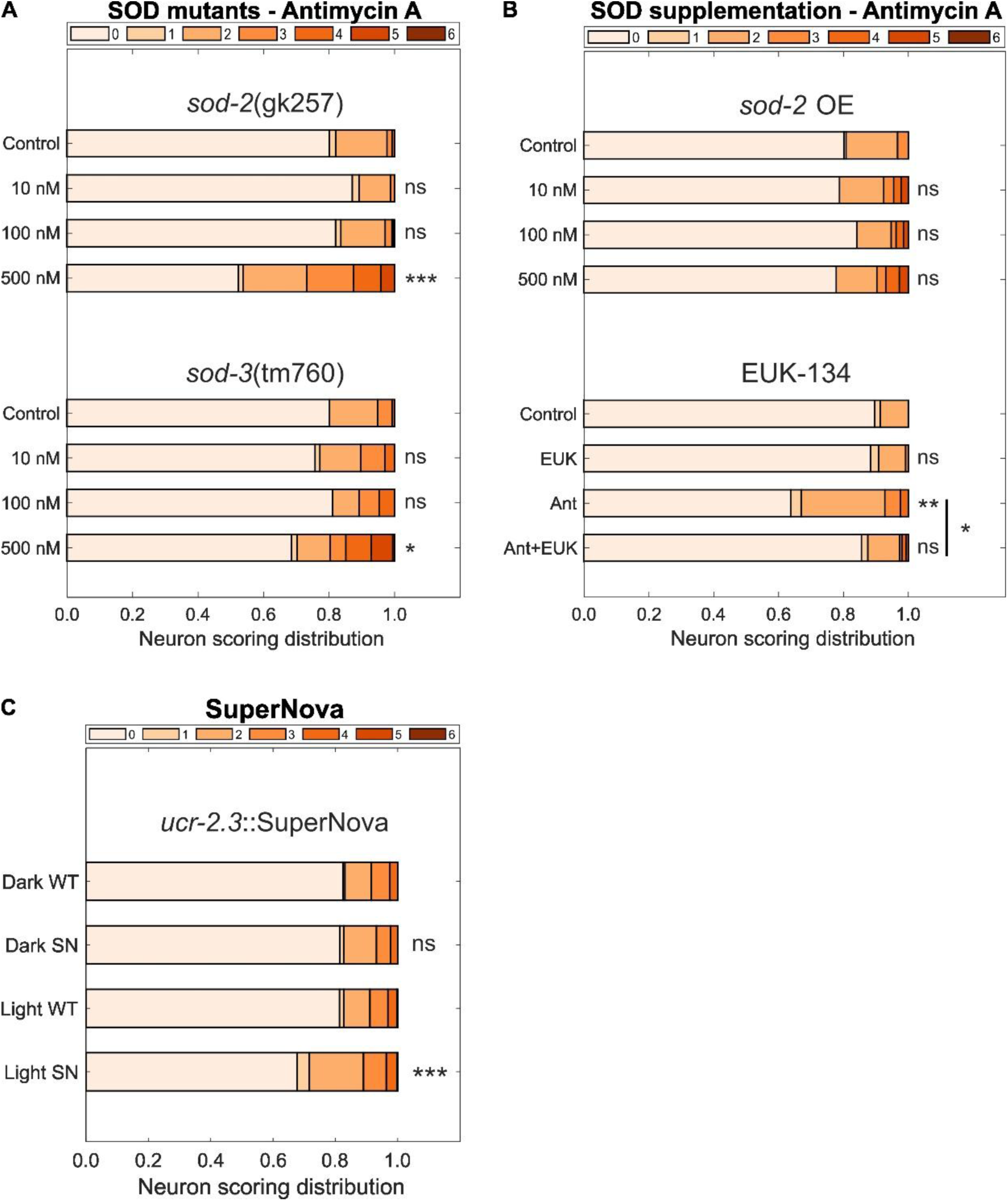
Mitochondrial superoxide anion regulates neurotoxicity linked to complex III inhibition. (**A**) Neuronal damage scoring distribution for cephalic dopaminergic neurons after developmental exposure to antimycin A in a *sod-2* and a *sod-3* mutant background. (**B**) Neuronal damage scoring distribution for cephalic dopaminergic neurons after developmental exposure to antimycin A in a *sod-2* over expressing strain and 500 nM antimycin A in combination with 0.5 mM EUK-134. (**C**) Neuronal damage scoring distribution after optogenetic activation of complex III SuperNova during development, SuperNova worms compared to their respective controls. *N* = 3 biological replicates (5 biological replicates for *sod-2* mutant and SuperNova), *n* = 50 - 80 dendrites per treatment per replicate, chi-square test with Bonferroni post-hoc test, ns *P* > 0.05, \**P* < 0.05, \*\**P* < 0.01, \*\*\**P* < 0.001.

## Discussion

Exposure to chemicals present in the environment as pollutants is an important contributor to the onset of PD and other neurodegenerative diseases [4]. Many of these chemicals are also mitochondrial toxicants [7]. However, mitochondrial dysfunction can take many different forms [9–11]. We focused this work on inhibition of complex III of the mitochondrial ETC by two pesticides, antimycin A and pyraclostrobin. We demonstrated that developmental exposure to either chemical led to selective dopaminergic neurotoxicity in the model organism *C. elegans*. This was directly linked to production of ROS in the complex III site, especially superoxide anion. Furthermore, we recapitulated dopaminergic neuronal damage independently of chemical exposure by generating ROS at complex III using a SuperNova construct. These results indicate that developmental inhibition of mitochondrial complex III by environmentally relevant chemicals leads to ROS-mediated selective dopaminergic neurotoxicity in *C. elegans*.

Complex III inhibition, like complex I inhibition, can both deplete ATP and cause ROS generation, suggesting that complex III inhibition might, like complex I inhibition, cause dopaminergic neurotoxicity. We focused on inhibition of complex III using the piscicide antimycin A [22] and fungicide pyraclostrobin [23]. We found that both chemicals caused dopaminergic-selective neurotoxicity at concentrations low enough to not cause whole-organism toxicity. We note that we chose to work with the highest concentration of pyraclostrobin allowed by the limit of solubility in our hands, because even this concentration had little or no effect on nematode growth rate. This concentration (50 µM; ∼20 mg/L) is higher than that reported in surface waters of 0.239 µg/L or the predicted concentration of 150 µg/L based on label application rates [66,67]. It is unclear how much pyraclostrobin was present internally (i.e., what the actual target-site dose was). The *C. elegans* cuticle offers a significant barrier to uptake of many chemicals, reducing their impact [68]. Overall, our results show selective effects on dopaminergic neurons, in that for both chemicals, at concentrations causing little or no growth inhibition, neuronal damage was not observed in other types of neurons. This result is further supported by the fact that of the functional outcomes we examined, only BSR, a dopaminergic-linked behavior, was altered by complex III inhibition.

Furthermore, we found that superoxide anion generated in the complex III site is causative of dopaminergic neurotoxicity in *C. elegans*, and that interventions to offset superoxide anion production after chemical inhibition of complex III reduced or abrogated neurotoxicity. A critical role of superoxide anion production in dopaminergic neurotoxicity has also been demonstrated for complex I inhibition [12,18], raising the question of whether this will be a common feature of all chemicals that cause dopaminergic neurotoxicity via mitochondrial disruption.

However, there are clearly other factors involved in dopaminergic neurotoxicity, raising the question of how they interact. One such factor is α-synuclein aggregation, which has a large role in PD [32]. Interestingly, although overexpression of human SNCA caused very significant dopaminergic neuronal damage on its own, antimycin A and pyraclostrobin appeared to have roughly additive effects when α-synuclein was present. Although our experimental design does not formally test additivity, both factors have independent effects, alone and together. Similarly, PRKN and PINK1 are mitophagy regulators that are linked to PD epidemiologically and mechanistically, due to their role in removing damaged proteins and mitochondria [6,34]. We only found evidence for *pdr-1* but not for *pink-1* mutations to have an interaction with chemically induced dopaminergic neurotoxicity. Interestingly, worms having both α-synuclein production and either a *pdr-1* or *pink-1* mutant background showed a loss of the significant neurotoxic effect of complex III inhibition. A caveat to these results is that it is possible that the high baseline of neuronal damage caused by α-synuclein in our model caused an obscuring effect of any additional neurotoxicity. Notably, α-synuclein is reported to cause mitochondrial fragmentation in *C. elegans*, with PINK1 and PRKN co-expression rescuing this inhibition of mitochondrial fusion by α-synuclein [69]. This led us to hypothesize a synergistic worsening of the neuronal damage of these combined backgrounds, but our results do not support this. We previously reported that *pdr-1* sensitizes *C. elegans* to dopaminergic neurodegeneration driven by 6-OHDA exposure, and *pink-1* had a protective effect under the same conditions [70]. This coincides with our observations of dopaminergic neurotoxicity for both backgrounds, with *pink-1* worms having less deviation from the control than those with a *pdr-1* mutation.

PD is the second most common neurodegenerative disease and the most common movement disorder, affecting approximately 10 million people worldwide globally as of 2021 [1]. This is likely to increase with a growing population of older people. Therefore, understanding the causes of PD to prevent cases is of critical importance. Exposure to environmental pollutants such as pesticides and heavy metals, have been linked to the onset of PD [5,71,72], but epidemiological evidence is scarce and causality often unclear. Mitochondrial toxicity is a mode of action for these toxicants, with strong evidence for complex I inhibition as causative of parkinsonism and dopaminergic neurodegeneration [12,13,44]. A recent screening of various pesticides targeting complexes I, II, and III using human cell culture models, found evidence that inhibition of both complex I and III led to dopaminergic neurotoxicity [15]. We are not aware of other studies testing selective dopaminergic neuron sensitivity to complex III inhibition. Our work provides robust confirmation that 1) complex III inhibition causes dopaminergic neurotoxicity *in vivo*, and 2) that this is directly related to generation of mitochondrial ROS in complex III, providing mechanistic insights on this specific mode of mitochondrial dysfunction. This is important from a regulatory perspective because complex III inhibitors are common in nature, frequently used as pesticides, and increasingly pursued for drug development [73]. Importantly, despite differences between the *C. elegans* model and vertebrate nervous systems, such as the absence of a blood-brain barrier or myelination, the scored morphological changes such as beading and breaks have been linked to mammalian neuronal phenotypes like neuronal beading on dendrites and age-dependent loss of dendritic branches respectively [74,75]. However, additional work in longer-lived models and epidemiological studies focusing specifically on complex III inhibitors will be important to assess the importance of complex III inhibition to PD or parkinsonism in people. From a mechanistic perspective, the knowledge that mitochondrial ROS, in particular superoxide anion, is at the center of dopaminergic dysfunction may be helpful in the development of preventive and therapeutic approaches.

## Materials and Methods

### *C. elegans* maintenance

*C. elegans* strains. Strains BY200 (*vtIs1*[p*dat-1*::GFP]), DA1240 (*adIs1240*[*eat-4*::sGFP+*lin-15*(+)]), LX929 (*vsIs48*[*unc-17*::GFP]), CZ1632 (*juIs76*[*unc-25*p::GFP+*lin-15*(+)]), GR1366 (*mgIs42*[*tph-1*::GFP+*rol-6*(su1006)]), VT1485 (*maIs188*[*mir-228*p::GFP+*unc-119*(+)], CB1112 (*cat-2*(e1112) II), CB1141 (*cat-4*(e1141) V), CB75 (*mec-2*(e75) X), CB1339 (*mec-4*(e1339) X), NM1657 (*unc-10*(md1117) X), VC223 (*tom-1*(ok285) I), ERS1 (*eraIs1*[*dat-1p*::mCherry, *dat-1p*::hSNCA::Venus]), ERS44 (*eraIs1*[*dat-1p*::mCherry, *dat-1p*::hSNCA::Venus]; *pdr-1*(gk448)), ERS49 (*eraIs1*[*dat-1p*::mCherry, *dat-1p*::hSNCA::Venus]; *pink-1(tm1779)*), UA226 (*pink-1*(tm1779); *vtIs1*[p*dat-1*::GFP]), UA227 (*pdr-1*(tm598); *vtIs1*[p*dat-1*::GFP]), PHX2867 (p*dat-1*::MLS::roGFP), PHX2923 (p*dat-1*::PercevalHR), GA184 (*sod-2*(gk257) I), GA186 (*sod-3*(tm760) X), GA805 (*wuIs156*[*sod-2*(genomic) + *rol-6*(su1006)], and APW125 (*jbm21*[*ucr2.3*::link::SuperNova] III) were maintained at 20 °C on K-agar plates seeded with OP50 *E. coli* [76]. Worms were fed OP50 *E. coli* in experiments requiring culture on solid medium because of the limited bacterial lawn obtained with OP50, allowing easier observation of worms [77]. HB101 *E. coli* was used in experiments requiring liquid culture because it is less prone to forming bacterial clumps, making it easier to eat for the worms [78]. Strain BY200 was a gift from Michael Aschner; strain APW125 was a gift from Andrew Wojtovich; strains ERS1, ERS44, and ERS49 were a gift from Roman Vozdek; and other strains were obtained from the Caenorhabditis Genetics Center (Table S1).

Generation of crossed strains. BY200 male worms were generated by exposing L4 stage worms to 33 °C for 4 hours. After 2-3 days, the offspring were screened for males and transferred to new plates containing one hermaphrodite and three males each. This step was necessary to increase the number of available males in the next generation. After 2-3 days, new plates were prepared containing one hermaphrodite of either of the GA184, GA185, GA805 or APW125 strains and three BY200 second-generation males. After 2-3 days, putative homozygous crosses were first verified for the visible BY200 dopaminergic neuron fluorescence. Putative homozygous crosses with strain GA805 were further screened for the *rol* phenotype. Putative homozygous crosses with strains GA184, GA185, and APW125 were verified by PCR genotyping (Table S2).

Age-synchronization by bleaching treatment. For experiments requiring age-synchronization, a non-starved population of day 1-2 adults was collected in a 15 mL tube from K-agar plates by washing with K-medium [76]. Worms were allowed to settle for 2-3 minutes, and the supernatant discarded. Worms were then treated with 5 mL of K-medium containing final concentrations of 0.4 N sodium hydroxide and 20% v/v sodium hypochlorite for eight minutes. The bleaching reaction was quenched by raising the volume to 15 mL with K-medium, centrifuged at 2200 RCF for 2 minutes, and the supernatant discarded. This washing last step was repeated two more times to recover the embryos for exposure experiments. We tested whether this bleaching treatment would affect dopaminergic neuronal damage after the worms reached the L4 larval stage. We found no significant effect on dopaminergic neurotoxicity by this treatment compared to worms that were age-synchronized by adult egg-laying (eggs laid for one hour, and then adults removed) or by egg isolation via bleaching followed by L1 synchronization by overnight hatch in liquid without food (Fig. S8).

### Chemical exposure design

Developmental exposure to chemicals. Embryos generated by bleaching treatment were counted on a stereo microscope by transferring 10 µL of K-medium containing embryos to a glass slide. K-medium was added or subtracted until reaching a concentration of 10 embryos/µL. Approximately 100 embryos in 10 µL of K-medium were transferred to each well of a 24-well plate and the volume was increased to a total of 500 µL per well of complete K-medium (0.5 mL of 10 mg/mL cholesterol, 3 mL of 1 M calcium chloride, 3 mL of 1 M magnesium sulfate per 1 L K-medium) containing HB101 *E. coli* bacterial food at a final concentration of OD 2.00, and either 0, 10, 100 or 500 nM of antimycin A (Sigma-Aldrich) or 0, 10, or 50 µM of pyraclostrobin (Sigma-Aldrich) final concentrations. All wells also contained a final concentration of 1% v/v DMSO as vehicle. The 24-well plate was put on a shaker at 20 °C for 52-54 hours to allow worms to reach the L4 larval stage. Afterward, liquid and worms from each well were collected in separate 15 mL tubes, the total volume raised to 10 mL with K-medium, and the worms were allowed to settle. After 2-3 minutes, the supernatant was discarded, and this washing step was repeated two more times. At this point worms were ready for endpoint measurement.

Determination of experimental concentrations. Embryos were treated as described in the preceding paragraph, but with additional concentrations tested during development; 0, 10, 25, 50, 100, or 500 nM antimycin A, or 0, 10, 25, or 50 µM pyraclostrobin final concentrations. After 52-54 hours worms were collected and transferred to unseeded K-agar plates, allowing the worms to disperse on the plate for 5 minutes. Plates containing worms were mounted and imaged using a Keyence BZ-X710 microscope using a Nikon 4X objective. Images were analyzed using the WormSizer add-in in ImageJ following their published protocol to determine worm length of a minimum of 50 individuals per treatment [79]. These length values were then used to construct a dose response curve using AAT Bioquest EC_50_ Calculator [24] to determine EC_10_ and EC_25_ concentrations.

6-hydroxydopamine exposure. Worms that were exposed to 6-OHDA as a secondary challenge or as a positive control for roGFP and PercevalHR assays were collected and washed as described in the preceding paragraphs. These worms were then transferred to 24-well plates and well volumes were raised to a total of 500 µL per well of complete K-medium containing 50 mM of 6-hydroxydopamine (Sigma-Aldrich) and 10 mM of L-ascorbic acid (Sigma-Aldrich) final concentrations. Control wells contained 500 µL per well of complete K-medium with 10 mM of L-ascorbic acid. Worms were exposed for one hour and then collected in 15 mL tubes and washed three times as described above. Microscopy was performed immediately after.

Acute antimycin A exposure. Embryos generated by bleaching treatment were counted on a stereo microscope by transferring 10 µL of K-medium containing embryos to a glass slide. Approximately 100 embryos in K-medium were transferred to K-agar plates and allowed to reach the L4 larval stage. After 52-54 hours, worms were collected and transferred to 24-well plates and volumes raised to a total of 500 µL per well of complete K-medium containing 0, or 10 µM of antimycin A final concentrations for 2.5 hours. All wells also contained a final concentration of 1% v/v DMSO as vehicle. Worms were collected in 15 mL tubes after 2.5 hours and washed as described above. Microscopy was performed immediately after.

### Quantification of neuronal damage

Microscopy imaging. L4 larval stage worms that previously went through chemical exposure were transferred to 5 µL of 100 mM sodium azide solution on a 2% agarose pad on top of a glass slide. Worms were paralyzed by the sodium azide after one minute and covered with a coverslip. The glass slide was mounted on a Keyence BZ-X710 microscope equipped with a Keyence BZ-X700E metal halide light-source. Z-stacks of individual worms’ heads were acquired using a Nikon 40X objective and a Chroma GFP filter cube with 100 milliseconds of exposure, microscope objective’s pitch of 0.5 µm, and 3×3 binning.

Late life scoring. L4 larval stage worms that went through chemical exposure during development were collected in 15 mL tubes and raised to 15 mL with K-medium. Worms were allowed to settle for 2-3 minutes and the supernatant discarded. This washing step was repeated two more times. Worms were transferred to K-agar plates containing OP50 *E. coli* lawns. Worm collection and washing was repeated every 1-2 days to allow for transfer to fresh plates while eliminating embryos and larvae. Microscopy was performed 6 days after exposure.

Neuronal damage quantification. Maximum projections of the z-stacks based on maximum intensity were generated, and each dendrite of the cephalic neurons was scored as previously described [21,26]. For dopaminergic neurons: 0 – no damage, 1 – irregular (curves), 2 – less than 5 blebs, 3 – 5 to 10 blebs, 4 – more than 10 blebs and/or breaks, 5 – breaks, 25 to 75% dendrite loss, and 6 – breaks, more than 75% dendrite loss. For cholinergic neurons: 0 – no damage, 1 – irregular (curves), 2 – 1 to 10 blebs, and 3 – more than 10 blebs and/or presence of breaks. For glutamatergic neurons: 0 – no damage, 1 – 1 to 5 blebs, 2 – 6 to 10 blebs, and 3 – more than 10 blebs and/or presence of breaks. For serotonergic neurons, GABAergic neurons and glial cells: 0 – no damage, and 1 – any type of damage (Fig. S2). Scoring was performed blinded by using the software Blinder following their published protocol [80].

### Behavioral assays

Basal slowing response. Assay plates were prepared by spreading OP50 *E. coli* in the shape of a ring on K-agar plates and incubated overnight. Worms that previously went through chemical exposure were washed and transferred to the center of either an assay plate or a plate with no bacteria. Excess liquid was absorbed using a Kimwipe to minimize the time required for drying. After 5 minutes, plates were mounted on a Keyence BZ-X710 microscope, and 20 second videos were acquired until a minimum of 10 individual worms were imaged. These videos were used to count the number of body bends on plates with and without bacterial food. Strains CB1112 and CB1141 were used as positive controls.

Sensitivity to soft touch. Worms that previously went through chemical exposure were washed and transferred to unseeded K-agar plates. Plates with worms were allowed to dry for 5 minutes. Worms were tested by stroking their heads and tails consecutively with an eyelash mounted on a Pasteur pipette under a Leica MZ75 stereo microscope. Sensitive worms started backwards movement after touching their heads and stopped or started forward locomotion after touching their tails. This process was repeated 5 times per worm for a minimum of 20 worms per plate, counting the number of times that a worm showed response to soft touch. Worms that did not start backwards motion after the initial head touch three consecutive times were also counted as not responsive for that try. Strains CB75 and CB1339 were used as positive controls.

Aldicarb-induced paralysis. Assay plates containing 1 mM aldicarb (Sigma-Aldrich) were prepared one day before the assay and stored at 4 °C. Aldicarb plates were allowed to reach room temperature before starting the assay. A drop of 2 μL of HB101 *E. coli* was added to the center of the plate and allowed to dry for 10 minutes. 20 to 30 worms that previously went through chemical exposure were washed and transferred to the bacterial spot on the plate. Paralysis was assessed every 30 minutes for 6 hours. A paralyzed worm was defined as not showing movement after prodding with an aluminum wire. Strains NM1657 and VC223 were used as positive controls.

### Quantification of glutathione redox tone and ATP:ADP ratio in dopaminergic neurons

Worms of strains PHX2867 and PHX2923, expressing mitochondrial-localized roGFP and PercevalHR respectively under the *dat-1* promoter [40], were used to quantify the ATP:ADP ratio and glutathione redox tone in dopaminergic neurons. Worms that previously went through chemical exposure (for 18, 24, or 52-54 hours) were transferred to 5 µL of K-medium on an 8% agarose pad on top of a glass slide and covered with a coverslip. No paralytic chemicals were used because many interfere with these parameters [81]. The glass slide was mounted on a Keyence BZ-X710 microscope equipped with a Keyence BZ-X700E metal halide light-source. Two images per head of an individual worm, focused on the cell bodies of the cephalic dopaminergic neurons, were acquired using a Nikon 40X objective, 25 milliseconds of exposure, and 3×3 binning. The first image was acquired using a 470ex/520em filter, and the second image with a 405ex/520em filter. A custom-made MATLAB script was used to perform feature-based segmentation of the cell bodies in each image, background subtraction, and to determine their mean intensity values. The script then calculated the mean intensity ratios of 405/470 excitation for mitochondrial roGFP as oxidized to reduced glutathione, and the mean intensity ratios of 470/405 excitation for PercevalHR as ATP to ADP ratio.

### Chemical rescue experiments

Embryos generated by bleaching treatment were counted on a stereo microscope by transferring 10 µL of K-medium containing embryos to a glass slide. K-medium was added or subtracted until reaching a concentration of 10 embryos/µL. Approximately 100 embryos in 10 µL of K-medium were transferred to each well of a 24-well plate and raised to a total volume of 500 µL per well of complete K-medium containing HB101 *E. coli* bacterial food at a final concentration of OD 2.00. For each replicate, one well contained no additional chemicals to serve as control, one well contained only the rescue chemical, one well contained 500 nM antimycin A, and one well contained 500 nM antimycin A and the rescue chemical. The rescue chemicals final concentrations were 2.5 mM N-acetylcysteine (Sigma-Aldrich), 100 µM S3QEL-2 (Sigma-Aldrich), 25 mM dichloroacetate (Sigma-Aldrich), 10 µg/mL febuxostat (Sigma-Aldrich), 10 µM myxothiazol (Sigma-Aldrich), or 0.5 mM EUK-134 (Sigma-Aldrich). All wells also contained a final concentration of 1% v/v DMSO as vehicle. The 24-well plate was put on a shaker at 20 °C for 52-54 hours to allow worms to reach the L4 larval stage. Afterward, liquid and worms from each well were collected in separate 15 mL tubes, raised to 10 mL with K-medium, and the worms were allowed to settle. After 2-3 minutes, the supernatant was discarded, and this washing step repeated two more times. At this point worms were ready for endpoint measurement.

### Optogenetic induction of SuperNova

Embryos generated by bleaching treatment were counted on a stereo microscope by transferring 10 µL of K-medium containing embryos to a glass slide. Approximately 100 embryos in K-medium were transferred to K-agar plates. SuperNova activation was conducted using the AMUZA 590 nm LED array for Multiwell Plates [12,63]. K-agar plates containing embryos were placed approximately 2 cm above the surface of the LED array location to enable air flow and to keep temperature in the incubator at 20 °C. Worms were under continuous light exposure of 0.3 mW/mm^2^ and allowed to reach the L4 larval stage. Dark controls were placed within the same incubator, in a cardboard box with an aluminum foil lid to prevent light penetration. After 52-54 hours, K-agar plates containing worms were retired from the incubator and microscopy was performed.

### Quantification of oxygen consumption rate (OCR)

We used a Seahorse XFe96 Extracellular Flux Analyzer (Agilent) to quantify OCR in L4 worms that previously went through chemical exposure following published protocols [35]. Briefly, worms were transferred to a Seahorse microplate at a concentration of 20 worms per well. Basal OCR measurements were taken before injection of either 25 μM final concentration of carbonyl cyanide 4-(trifluoromethoxy) phenylhydrazone (FCCP) (Sigma-Aldrich) to measure maximal oxygen consumption or 20 μM final concentration of N,N-dicyclohexylcarbodiimide (DCCD) (Sigma-Aldrich), necessary to quantify basal mitochondrial OCR. After the injection of either FCCP or DCCD, 14 measurements of OCR were performed before a final injection of 10 mM final concentration of sodium azide (Sigma-Aldrich) to inhibit mitochondrial respiration and quantify non-mitochondrial OCR. Final parameters calculated include basal OCR, basal mitochondrial OCR, maximal OCR, spare capacity, proton leak, and non-mitochondrial OCR. OCR experiments included 5 wells per treatment group (technical replicates) for each biological replicate. OCR calculated values were normalized to worm count and worm volume. Average worm volume was calculated by acquiring images of a subset of 50-100 worms per group and using the ImageJ plugin WormSizer following their published protocol [79].

## Statistical analysis

OASIS2 was used for analysis of paralysis induced by aldicarb [82]. MATLAB R2024b (MATLAB 24.2) was used for all other statistical testing and graph generation. Statistical tests and sample size are described in their corresponding figure legends.

## Data availability

Data for this article is available at Duke University Digital Repository at https://doi.org/10.7924/r4bc47n2v.

## Author contributions

JH: conceptualization, methodology, software, formal analysis, investigation, visualization, writing – original draft, writing – review & editing. AW: formal analysis, investigation. JJ: formal analysis, investigation. JNM: conceptualization, methodology, supervision, project administration, funding acquisition, writing – original draft, writing – review & editing.

