## Supplemental data for "Inhibition of mitochondrial complex III causes selective dopaminergic neurotoxicity by redox stress in *Caenorhabditis elegans*"

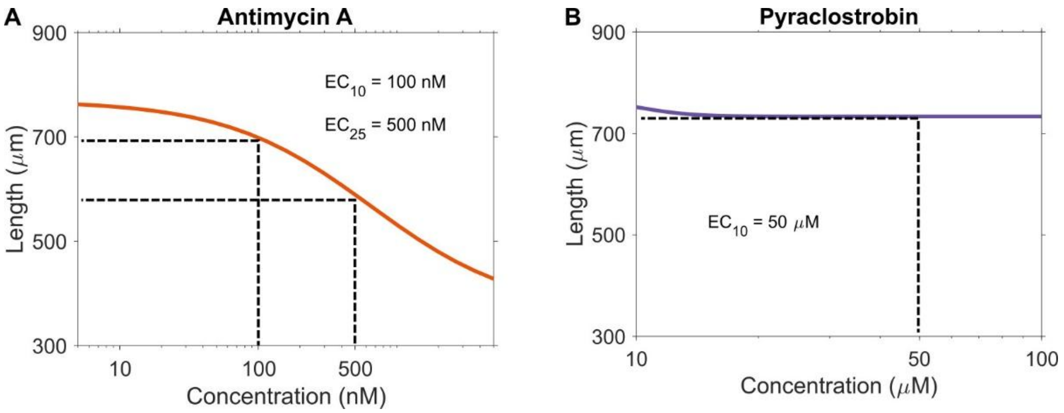


**Figure S1: Dose response curves.** Length-based response curves for (**A**) antimycin A and (**B**) pyraclostrobin. $EC_{10}$ = 100 nM and $EC_{25}$ = 500 nM for antimycin A, $EC_{10}$ = 50 µM for pyraclostrobin.

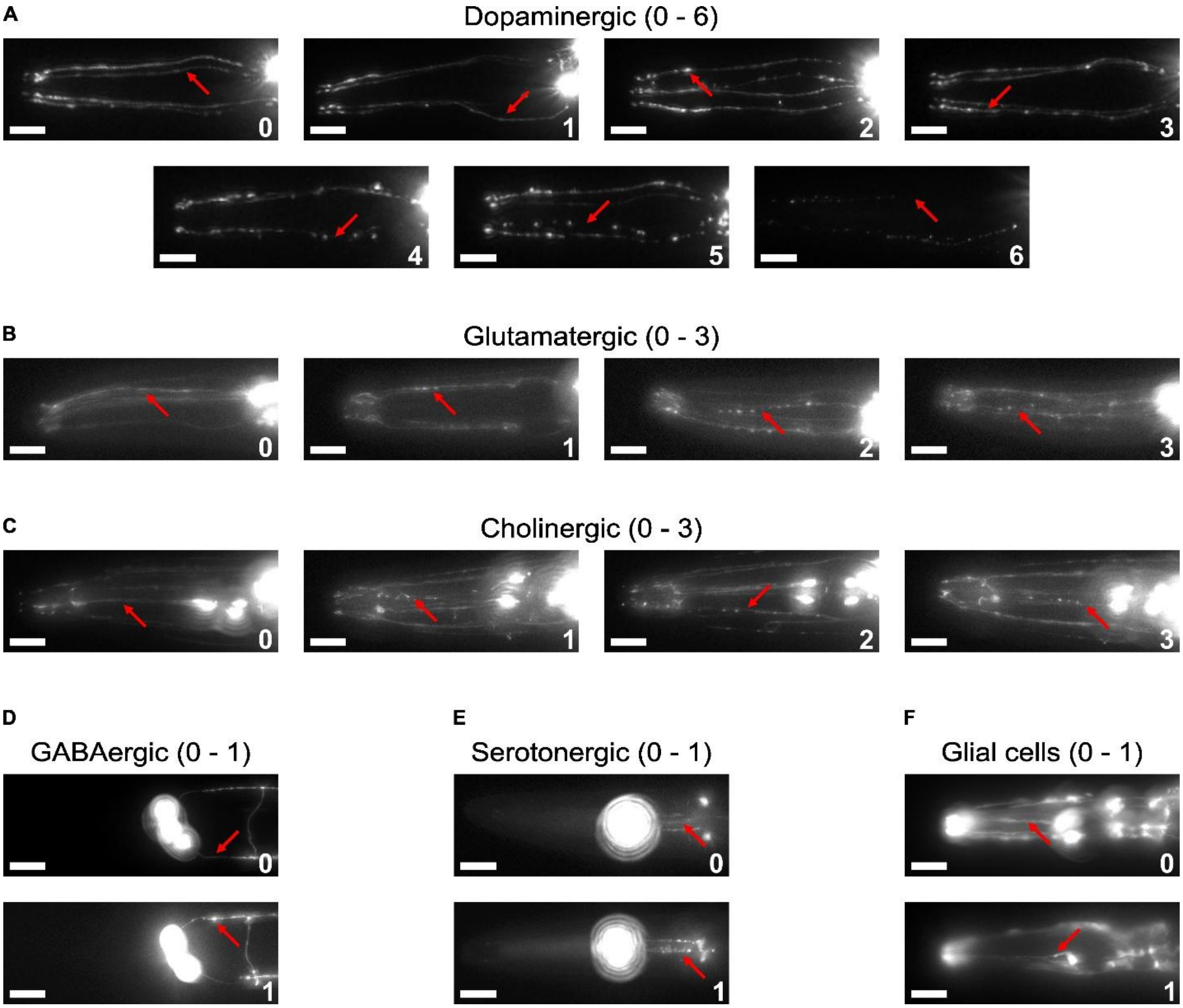


**Figure S2. Scoring of neuronal damage.** Representative images for each scoring category for (**A**) dopaminergic, (**B**) glutamatergic, (**C**) cholinergic, (**D**) GABAergic, and (**E**) serotonergic neurons, and (**F**) glial cells located in the head of *C. elegans*; red arrows indicate the scored neurite. Scale bars 20 µm.

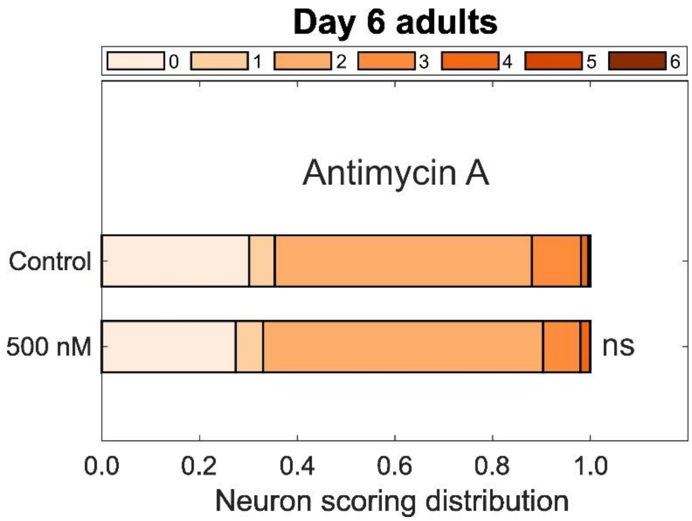


**Figure S3: Neuronal damage in late life.** Neuronal damage scoring distribution for cephalic dopaminergic neurons of day 6 adults that were exposed to 500 nM antimycin A during development. $N = 3$ biological replicates, $n = 60\text{-}80$ dendrites per treatment per replicate, chi-square test with Bonferroni post-hoc test, ns $P > 0.05$.

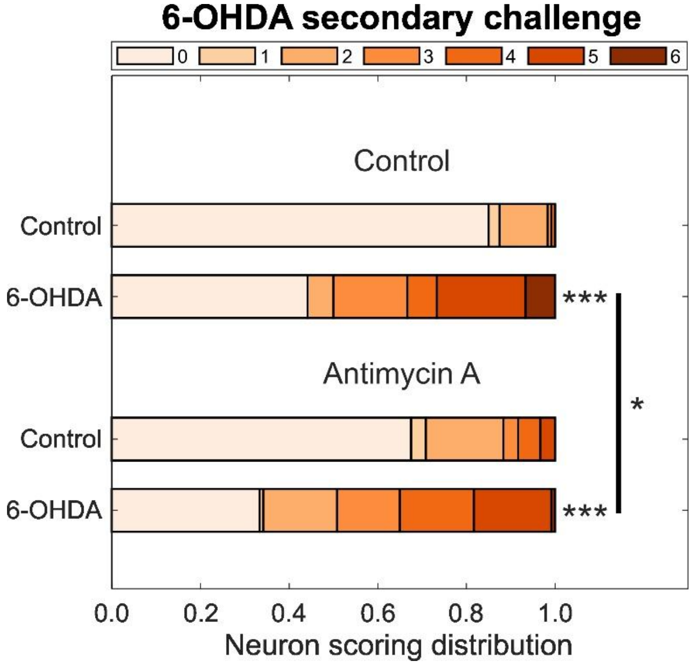


**Figure S4. Neuronal damage after 6-OHDA secondary challenge.** Neuronal damage scoring distribution for cephalic dopaminergic neurons after one hour challenge with 50 mM 6-OHDA following developmental exposure to 500 nM antimycin A. $N = 3$ biological replicates, $n = 40$ dendrites per treatment per replicate, chi-square test with Bonferroni post-hoc test, $^{*}P < 0.05$, $^{***}P < 0.001$.

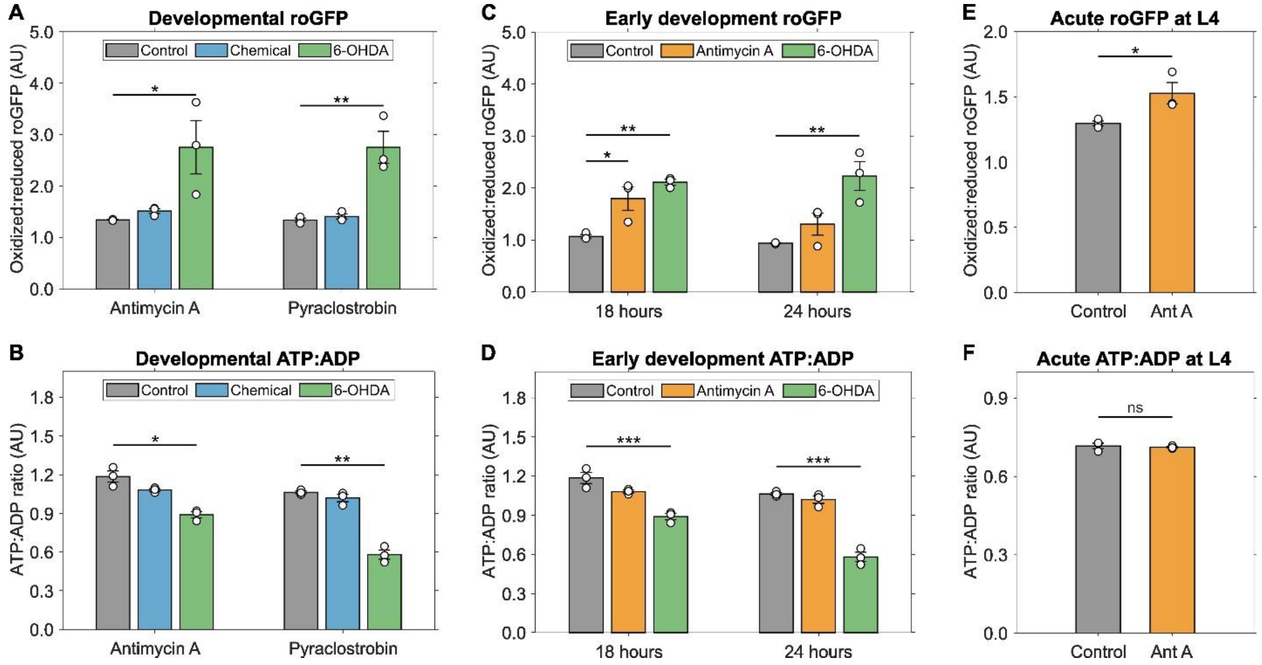


**Figure S5: Complex III inhibition alters dopaminergic redox but not bioenergetic state during early development, not normalized.** (**A**) Oxidized to reduced roGFP ratio in cephalic dopaminergic neurons of *C. elegans* developmentally exposed to 500 nM antimycin A or 50 µM pyraclostrobin; (**B**) 500 nM antimycin A after 18 or 24 hours of exposure. (**C**) ATP to ADP ratio as measured using the PercevalHR reporter in cephalic dopaminergic neurons of *C. elegans* developmentally exposed to 500 nM antimycin A or 50 µM pyraclostrobin; (**D**) 500 nM antimycin A after 18 or 24 hours of exposure. 6-OHDA included as positive control. (**E**) Oxidized to reduced roGFP ratio and (**F**) ATP to ADP ratio as measured using the PercevalHR reporter in cephalic dopaminergic neurons of *C. elegans* exposed to 10 µM antimycin A for 2.5 hours at the L4 larval stage. $N = 3$ biological replicates, $n = 20$-40 neurons per treatment per replicate, one-way ANOVA test with Dunnett's post-hoc test, ns $P > 0.05$, *$P < 0.05$, **$P < 0.01$, ***$P < 0.001$.

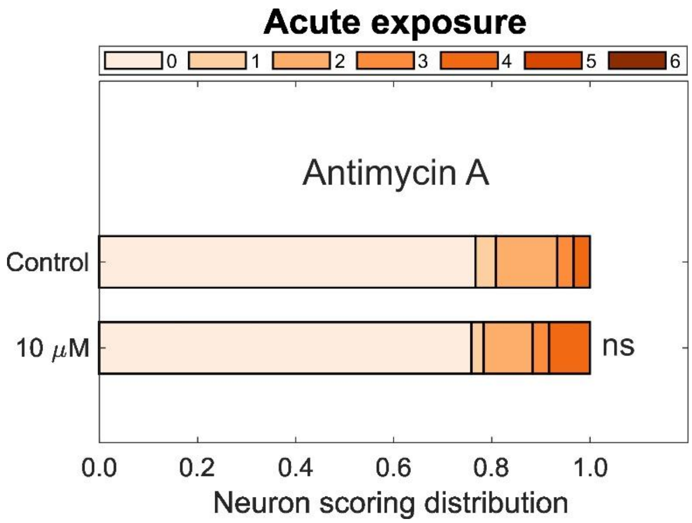


**Figure S6. Acute exposure to antimycin A does not cause dopaminergic neuronal damage.** Neuronal damage scoring distribution for cephalic dopaminergic neurons after two and a half hours exposure to 10 μM antimycin A at the L4 larval stage. $N = 3$ biological replicates, $n = 40$ dendrites per treatment per replicate, chi-square test with Bonferroni post-hoc test, ns $P > 0.05$.

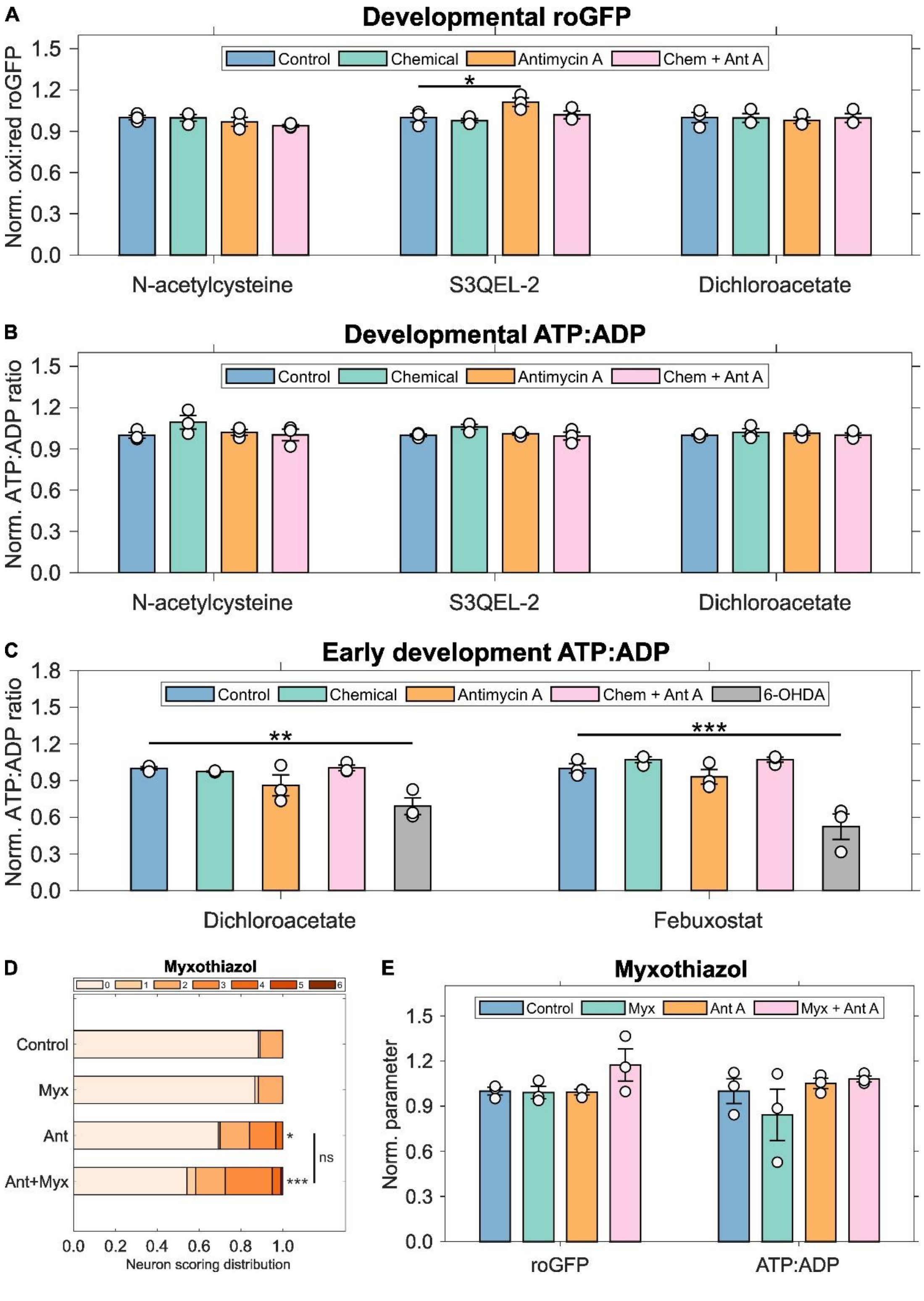


**Figure S7. Dopaminergic redox state and ATP levels after chemical rescue.** (**A**) Oxidized to reduced roGFP ratio and (**B**) ATP to ADP ratio measured using the PercevalHR reporter in cephalic dopaminergic neurons of *C. elegans* developmentally exposed to 500 nM antimycin A with 2.5 mM N-acetylcysteine, 100 µM S3QEL-2, and 25 mM dichloroacetate. $N = 3$ biological replicates, $n = 20$ neurons per treatment per replicate, one-way ANOVA test with Dunnett's post-hoc test. (**C**) ATP to ADP ratio measured using the PercevalHR reporter in cephalic dopaminergic neurons of *C. elegans* measured after 18 hours of exposure to 500 nM antimycin A with 25 mM dichloroacetate, and 10 µM febuxostat. 6-OHDA included as positive control. $N = 3$ biological replicates, $n = 40$ neurons per treatment per replicate, one-way ANOVA test with Dunnett's post-hoc test. (**D**) Neuronal damage scoring distribution for cephalic dopaminergic neurons after exposure to 500 nM antimycin A and 10 µM myxothiazol. $N = 3$ biological replicates, $n = 40$-60 dendrites per treatment per replicate, chi-square test with Bonferroni post-hoc test. (**E**) Oxidized to reduced roGFP ratio and ATP to ADP ratio measured using the PercevalHR reporter in cephalic dopaminergic neurons of *C. elegans* developmentally exposed to 500 nM antimycin A and 10 µM myxothiazol. $N = 3$ biological replicates, $n = 20$ neurons per treatment per replicate, one-way ANOVA test with Dunnett's post-hoc test.; ns $P > 0.05$, *$P < 0.05$, **$P < 0.01$, ***$P <$ 0.001.

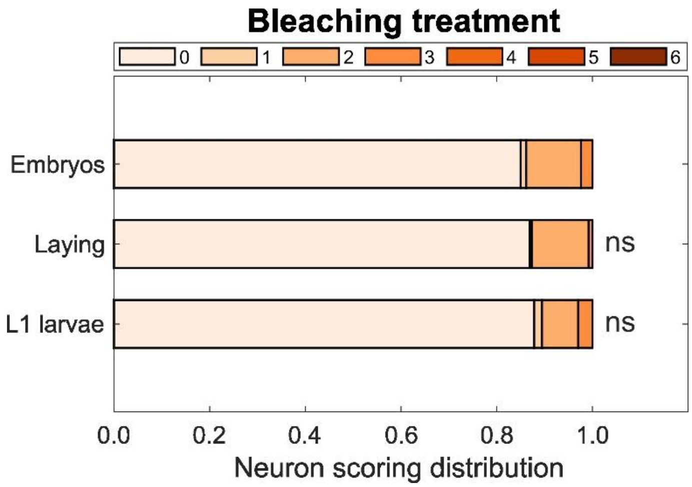


**Figure S8. Neuronal damage after bleaching treatment.** Neuronal damage scoring distribution for cephalic dopaminergic neurons measured at the L4 larval stage of *C. elegans* populations age-synchronized by allowing adults to lay eggs, by egg recovery from gravid adults after bleaching treatment, or gravid adult bleaching treatment followed with overnight egg hatching to produce L1 larval stage worms. $N = 3$ biological replicates, $n = 80$ dendrites per treatment per replicate, chi-square test with Bonferroni post-hoc test, ns $P > 0.05$.

**Table S1: List of *C. elegans* strains.**

| **Strain name** | **Genotype** | **Description** |
|---|---|---|
| BY200 | *vtIs1*[p*dat-1*::GFP] | Dopaminergic reporter |
| DA1240 | *adIs1240*[*eat-4*::sGFP+*lin-15*(+)] | Glutamatergic reporter |
| LX929 | *vsIs48*[*unc-17*::GFP] | Cholinergic reporter |
| CZ1632 | *juIs76*[*unc-25*p::GFP+*lin-15*(+)] | GABAergic reporter |
| GR1366 | *mgIs42*[*tph-1*::GFP+*rol-6*(su1006)] | Serotonergic reporter |
| VT1485 | *maIs188*[*mir-228*p::GFP+*unc-119*(+) | Glial cell reporter |
| CB1112 | *cat-2*(e1112) II | BSR defective strain |
| CB1141 | *cat-4*(e1141) V | BSR defective strain |
| CB75 | *mec-2*(e75) X | Soft touch defective strain |
| CB1339 | *mec-4*(e1339) X | Soft touch defective strain |
| NM1657 | *unc-10*(md1117) X | Aldicarb resistant strain |
| VC223 | *tom-1*(ok285) I | Aldicarb sensitive strain |
| ERS1 | *eraIs1*[*dat-1p*::mCherry, *dat-1p*::hSNCA::Venus] | Dopaminergic α-synuclein |
| ERS44 | *eraIs1*[*dat-1p*::mCherry, *dat-1p*::hSNCA::Venus]; *pdr-1*(gk448) | Dopaminergic α-synuclein, *pdr-1* mutant |
| ERS49 | *eraIs1*[*dat-1p*::mCherry, *dat-1p*::hSNCA::Venus]; *pink-1(tm1779)* | Dopaminergic α-synuclein, *pink-1* mutant |
| UA226 | *pink-1*(tm1779); *vtIs1*[p*dat-1*::GFP] | Dopaminergic reporter, *pink-1* mutant |
| UA227 | *pdr-1*(tm598); *vtIs1*[p*dat-1*::GFP] | Dopaminergic reporter, *pdr-1* mutant |
| PHX2867 | p*dat-1*::MLS::roGFP | Dopaminergic roGFP reporter |
| PHX2923 | p*dat-1*::PercevalHR | Dopaminergic PercevalHR reporter |
| GA184 | *sod-2*(gk257) I | *sod-2* mutant |
| GA186 | *sod-3*(tm760) X | *sod-3* mutant |
| GA805 | *wuIs156*[*sod-2*(genomic) + *rol-6*(su1006) | *sod-2* over expressor |
| APW125 | *jbm21*[*ucr2.3*::link::SuperNova] III | Complex III SuperNova |
| BY200 x GA184 | *vtIs1*[p*dat-1*::GFP]; *sod-2*(gk257) I | Dopaminergic reporter, *sod-2* mutant |
| BY200 x GA186 | *vtIs1*[p*dat-1*::GFP]; *sod-3*(tm760) X | Dopaminergic reporter, *sod-3* mutant |
| BY200 x GA805 | *vtIs1*[p*dat-1*::GFP]; *wuIs156*[*sod-2*(genomic) + *rol-6*(su1006) | Dopaminergic reporter, *sod-2* over expressor |
| BY200 x APW125 | *vtIs1*[p*dat-1*::GFP]; *jbm21*[*ucr2.3*::link::SuperNova] III | Dopaminergic reporter, complex III SuperNova |

**Table S2: Primers used for PCR genotyping of *C. elegans* crosses.**

| Strain 1 | Strain 2 | Forward primer | Reverse primer |
|---|---|---|---|
| BY200 | GA184 | CAAGTCCAGTTGTTGCCTCA | CAAAAACCACTTTGCTCGGT |
| BY200 | GA186 | TGTTTACTTTGTTCTCGTGGGTT | CCTTCCAAATAGCATGGACATAG |
| BY200 | APW125 | AGTCGAGATTAGCCCTTGGT | AAGCTCCTCTCTGTCTTGGC |
